# Phylogenetic conflicts, combinability, and deep phylogenomics in plants

**DOI:** 10.1101/371930

**Authors:** Stephen A. Smith, Nathanael Walker-Hale, Joseph F. Walker, Joseph W. Brown

## Abstract

Studies have demonstrated that pervasive gene tree conflict underlies several important phylogenetic relationships where different species tree methods produce conflicting results. Here, we present a means of dissecting the phylogenetic signal for alternative resolutions within a dataset in order to resolve recalcitrant relationships and, importantly, identify what the dataset is unable to resolve. These procedures extend upon methods for isolating conflict and concordance involving specific candidate relationships and can be used to identify systematic error and disambiguate sources of conflict among species tree inference methods. We demonstrate these on a large phylogenomic plant dataset. Our results support the placement of *Amborella* as sister to the remaining extant angiosperms, Gnetales as sister to pines, and the monophyly of extant gymnosperms. Several other contentious relationships, including the resolution of relationships within the bryophytes and the eudicots, remain uncertain given the low number of supporting gene trees. To address whether concatenation of filtered genes amplified phylogenetic signal for relationships, we implemented a combinatorial heuristic to test combinability of genes. We found that nested conflicts limited the ability of data filtering methods to fully ameliorate conflicting signal amongst gene trees. These analyses confirmed that the underlying conflicting signal does not support broad concatenation of genes. Our approach provides a means of dissecting a specific dataset to address deep phylogenetic relationships while also identifying the inferential boundaries of the dataset.

## Introduction

Researchers have amassed large genomic and transcriptomic datasets meant to resolve fundamental phylogenetic relationships in plants (Wickett et al. 2014), animals (Jarvis et al. 2014; Dunn et al. 2008; Simion et al. 2017; Whelan et al. 2017), fungi (Shen et al. 2016), and bacteria (Ahrenfeldt et al. 2017). While the goals of these data collection efforts have been to increase the overall phylogenetic support, analyses have demonstrated that different datasets and analytical approaches often reconstruct strongly supported but conflicting relationships (Feuda et al. 2017; Walker et al. 2018; Shen, Hittinger, and Rokas 2017). Underlying these discordant results are strongly conflicting gene trees (Smith et al. 2015). In some cases, one or two “outlier” genes with large likelihood differences between alternative relationships can influence inferred species relationships (Shen, Hittinger, and Rokas 2017; Brown and Thomson 2017; Walker, Brown, and Smith 2018). Analyzing the behavior of each individual gene tree within a multi-gene dataset appears key to understanding the difficulties in resolving the most recalcitrant relationships in the tree of life.

Phylogenomic datasets are often analyzed as concatenated supermatrices or with coalescent gene-tree / species tree methods. Supermatrix methods were, in part, developed to amplify the strongest phylogenetic signal. However, it has long been understood that the “total evidence” paradigm (Kluge 1989), where the true history will ‘win out’ if enough data are collected, is untenable. Genes with real and conflicting histories are expected within datasets due to biological processes like hybridization and incomplete lineage sorting (ILS) (Maddison 1997) in addition to outlying genes and sites as mentioned above (Shen, Hittinger, and Rokas 2017; Brown and Thomson 2017; Walker, Brown, and Smith 2018). “Species tree” inference involves estimating species relationships using methods that account for gene tree conflict due to ILS (Edwards, Liu, and Pearl 2007; Liu et al. 2009; Edwards 2009; Edwards et al. 2016) and are commonly conducted alongside concatenated supermatrix analyses in phylogenomic studies. Differences in the results from these two approaches are often explained by the differences in assumptions each makes. The concatenated supermatrix allows for mixed molecular models and gene-specific branch lengths but assumes a single underlying tree topology shared by all genes. This procedure is known to perform poorly in the presence of extensive ILS. Coalescent approaches, depending on the implementation, may assume that all conflict is the result of ILS (but see Boussau et al. 2013 and Ané et al. 2006), that all genes evolved under selective neutrality and constant effective population size, that all genes contain enough information to properly resolve nodes, and that gene trees are estimated accurately (Springer and Gatesy 2016).

Supermatrix and coalescent methods perform well in many scenarios. However, when unresolved nodes or discordance between species trees remain after large data collection efforts, relatively few methods exist for dissecting the phylogenetic signal within datasets. For example, Bayesian methods have been developed that incorporate processes that lead to gene tree discordance in addition to ILS (Ané et al. 2006; Boussau et al. 2013). These methods are often computationally intractable for large datasets encompassing thousands of genes and may not handle systematic error well. Recently, network methods that scale to large datasets have been developed (Solís-Lemus 2016, Wen et al. 2018), but these do not allow for dissecting signal within datasets. Filtering approaches where subsets of genes are analyzed based on model similarity or the relationships displayed by the genes (Chen, Liang, and Zhang 2015; Shen et al. 2016; Smith, Brown, and Walker 2018), help to enable computational tractability and distill signal. For example, Chen, Liang, and Zhang (2015) filtered for hypothesis-specific genes in the phylogeny of jawed vertebrates using two methods: one where only gene trees capable of supporting one of three resolutions for a given relationship were included in the analysis, and another where only gene trees which agreed with a widely-accepted control locus were retained for the analysis. Researchers have also examined alternative phylogenetic hypotheses in order to isolate the supporting signal (Shen, Hittinger, and Rokas 2017; Brown and Thomson 2017; Walker, Brown, and Smith 2018).

In plants, several large data collection efforts aimed at resolving difficult nodes have found extensive conflicts (Smith et al. 2015; Walker et al. 2018, 2017; Wickett et al. 2014). Resolution of these clades is not only important for systematics, but crucial to an evolutionary understanding of key biological questions. For example, the relationships among the lineages of bryophytes (hornworts, liverworts, and mosses) remain unclear despite extensive data collection and analysis efforts (Wickett et al. 2014; Puttick et al. 2018). One of the most heavily debated lineages in plant phylogenetics is the monotypic *Amborella*, the conflicting placement of which alters our understanding of early flowering plant evolution. *Amborella* has been inferred as sister to Nymphaeales, as sister to all angiosperms, or as sister to the remaining Angiosperms excluding Nymphaeales (Xi et al. 2014). The placement of *Amborella*, along with other contentious relationships across land plants, would provide greater confidence in our understanding of the evolution of early reproductive ecology, the evolution of floral development, and the life history of early land plants (Feild et al. 2004; Sauquet et al. 2017).

We conducted a detailed analysis of conflict and signal across a phylogenomic dataset in hopes of presenting a computationally tractable and practical way to examine contentious relationships. We extended methods for examining phylogenetic alternatives and present an approach that can be widely applied to empirical datasets to determine the support, or lack thereof, for phylogenetic hypotheses. We applied these methods to a large plant genomic dataset (Wickett et al. 2014), and identified systematic error, nested conflicting relationships, and support for alternative resolutions. We explore a practical means to test the topological combinability of subsets of genes based on a combinatorial heuristic and information criteria statistics. By taking this broad, information-centric approach, we hope to shed more light on the evolution of plants and present a tractable approach for dissecting signal with broad applicability for phylogenomic datasets across the Tree of Life.

## Materials and Methods

### Datasets

We analyzed the Wickett et al. (2014) dataset of transcriptomes and genomes covering plants (available from http://datacommons.cyverse.org/browse/iplant/home/shared/onekp_pilot). There were several different filtering methods and approaches used in the original manuscript and, based on conversations with the corresponding author, we analyzed the filtered nucleotide dataset with third codon positions removed. These sites were removed because of problems with excessive variation and GC content that caused problems with the placement of the lycophytes (Wickett et al. 2014). This dataset consisted of 852 aligned genes. We did not conduct any other filtering or alteration of these data before conducting the analyses performed as part of this study.

### Phylogenetic analyses

We inferred gene trees for each of the 852 genes using IQ-TREE (v. 1.6.3; Nguyen et al. 2015). We used the GTR+G model of evolution and calculated maximum likelihood trees along with SH-aLRT values (Guindon et al. 2010). For all constrained analyses, we conducted additional maximum likelihood analyses with the same model of evolution but constrained on the relationship of interest, although the rest of the tree topology was free to vary. These constrained analyses were conducted with RAxML (v. 8.2.12; Stamatakis 2018).

### Conflict analyses

We conducted several different conflict analyses. First, we identified the congruent and conflicting branches between the maximum likelihood gene trees (ignoring branches that had less than 80% SH-aLRT; Guindon et al. 2010), and the concatenated maximum likelihood species tree from the original publication (Fig. 2; Wickett et al. 2014). These analyses were conducted using the program **`bp`** available from https://github.com/FePhyFoFum/gophy. We placed these conflicting and supporting statistics in a temporal context by calculating the divergence times of each split based on the TimeTree of Life (Hedges, Dudley, and Kumar 2006; Hedges et al. 2015). By examining the dominant conflicting alternatives, we established which constraints to implement and compare for further analyses. Because the gene regions contain partially overlapping taxa, automated discovery of all conflicting relationships concurrently can be challenging. To overcome these challenges, we examine each constraint individually.

To determine the difference in the log-likelihood (lnL) values among conflicting resolutions, we conducted constrained phylogenetic analyses (with parameters described in the *Phylogenetic analyses* section above) and compared the lnL values of the alternative resolutions. We examined those results that had a difference in the lnL of greater than 2, considering this difference as statistically significant (Edwards 1984). For each gene, we noted the resolution with the highest lnL and took the difference between that and the second-best resolution’s lnL (ΔlnL). For each resolution, we then summed this quantity across all genes supporting that resolution. We excluded genes that contained obvious errors (e.g., outgroup relationships falling within ingroups).

**Table 1.**
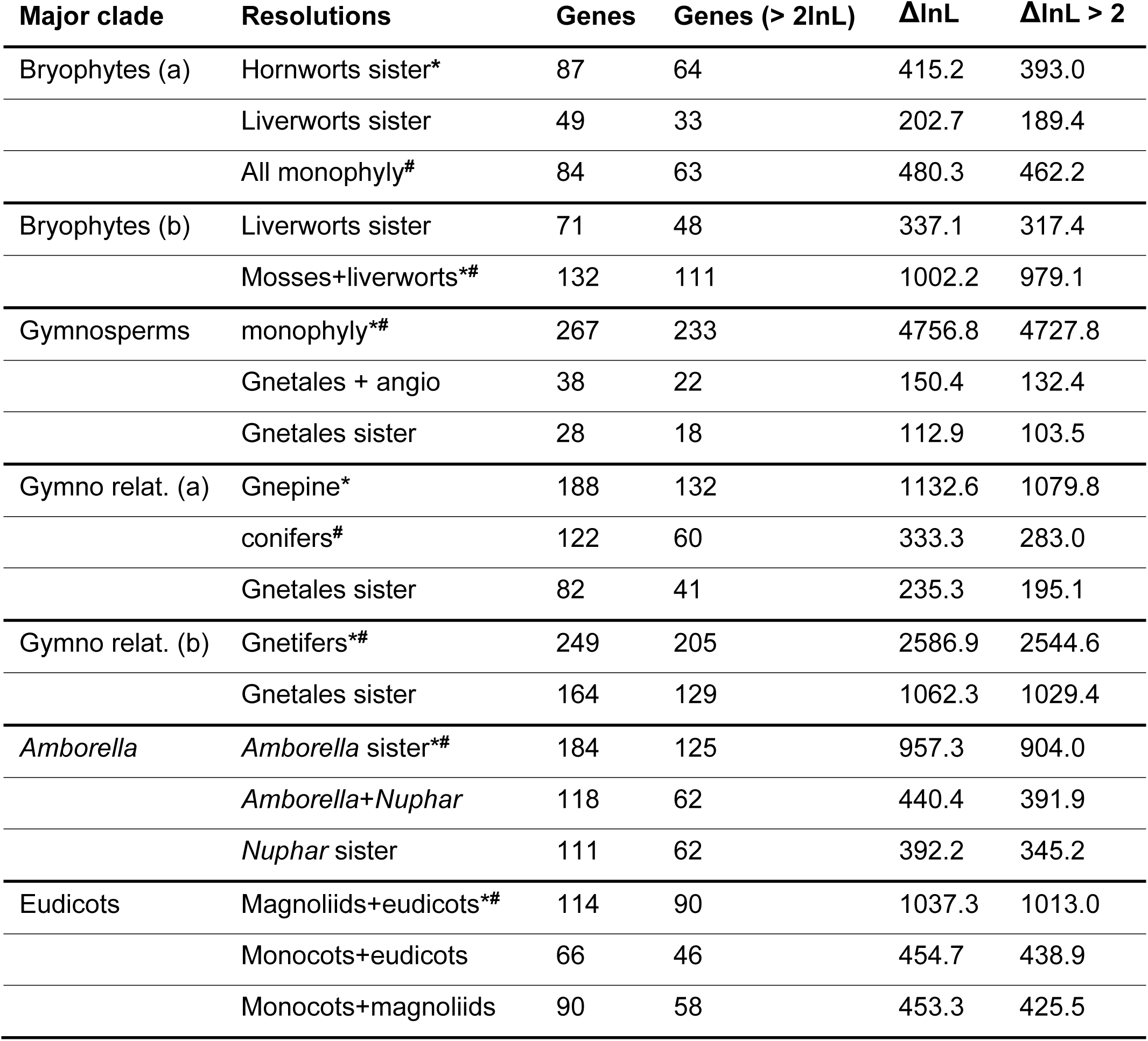
Comparison of the number of genes and the difference in the likelihood (ΔlnL) with relationships described in the Methods. *indicates relationships present in the ML tree and #indicates relationships present in the coalescent tree.

We examined five areas of plant evolution that highlight historical or current phylogenetic debates. First, we examined relationships involving Bryophyta, including hornworts sister to Embryophyta (i.e., monophyletic Embryophyta with the exclusion of hornworts), liverworts sister to Embryophyta (i.e. monophyletic Embryophyta with the exclusion of liverworts), mosses sister to liverworts (i.e. mosses and liverworts constrained to form a monophyletic group), and monophyletic bryophytes. Because mosses sister to liverworts does not conflict with hornworts sister to Embryophyta, we examined two sets of comparisons: (a) hornworts sister to Embryophyta, liverworts sister to Embryophyta, and monophyletic bryophytes, and (b) mosses sister to liverworts and liverworts sister to Embryophyta. Second, we examined alternative relationships regarding the gymnosperms (Acrogymnospermae), including monophyletic gymnosperms, Gnetales sister to angiosperms (i.e., monophyletic Gnetales and angiosperms), and Gnetales sister to angiosperms and gymnosperms (i.e., monophyletic angiosperms and gymnosperms with the exclusion of Gnetales). Third, within the gymnosperms we examined Gnepine (i.e., monophyletic Gnetales and Pinaceae), conifers (i.e., Pinaceae, Cupressaceae, Podocarpaceae, *Sciadopitys*), Gnetifers (i.e., monophyletic Gnetales and conifers), and Gnetales sister to the remaining gymnosperms (i.e., monophyletic conifers, cycas, and ginkgo). Because Gnetifers does not conflict with Gnepine, we examined two sets of comparisons: (a) Gnepine, conifers, and Gnetales sister to the remaining gymnosperms and (b) Gnetifers and Gnetales sister to the remaining gymnosperms. Note that conifers and Gnetales sister to the remaining gymnosperms also does not conflict but as our focal relationship for that comparison is Gnepine, they both conflict with the focal relationship and so are included. Fourth, we examined alternative placements of *Amborella*, including *Amborella* sister to angiosperms (i.e., monophyletic angiosperms with the exclusion of *Amborella*), *Amborella* sister to *Nuphar* (i.e., monophyletic *Amborella* and *Nuphar*), and *Nuphar* sister to angiosperms (i.e., monophyletic angiosperms with the exclusion of *Nuphar*). Finally, we examined alternative placements of eudicots and relatives including magnoliids as sister to eudicots (i.e., monophyletic magnoliids and eudicots), monocots sister to eudicots (i.e., monophyletic monocots and eudicots), and monocots sister to magnoliids (i.e., monophyletic magnoliids and monocots). In this case, *Sarcandra* was placed with eudicots given the placement in the tree in the original analyses and among the gene trees, though it was not required to be sister to the eudicots. These were not intended to be exhaustive and there may be additional reasonable hypotheses to examine. Instead, these were chosen to highlight the approaches discussed here.

We also examined nested conflicts. For the genes identified as supporting the dominant relationship of the eudicot lineages, we examined the distribution of conflict at other nodes in the tree. We first examined those genes that displayed both the eudicot lineages and the relationship of *Amborella* as sister to the rest of angiosperms. Then, of those genes, we determined which displayed the alternative gymnosperm relationships.

### Combinability test

To further examine whether there was support for concatenation of genes within the dataset after filtering for agreement in specific relationships, we developed and describe a simple procedure for testing the combinability within a dataset based on gene tree similarity and information criteria (Fig. 1). A typical concatenated phylogenetic analysis assumes that the entire alignment used to calculate the tree was generated with the same underlying topology. It follows that those genes that should be combined (i.e., concordant histories) will have more similar gene trees than those that should be considered separately (i.e., conflicting histories). To determine similarity between gene trees, we calculated the pairwise Robinson-Foulds (RF) distance (Robinson and Foulds 1981). We then constructed a graph where genes are nodes and edges are the weighted distances between gene trees based on RF. This can also be constructed with weighted RF. Then, beginning with the strongest edge, we tested for the combinability between the two connecting nodes. If they were combinable, based on the information criteria discussed below, we merged the nodes, along with the connecting edges for each.

**Figure 1.**
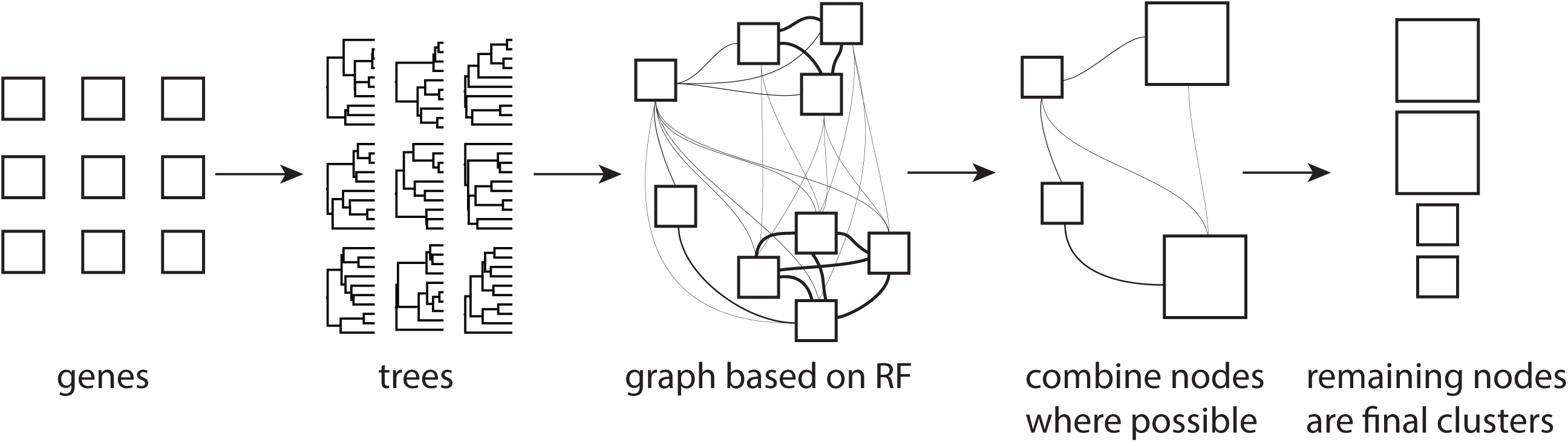
Procedure described in the methods section. Gene trees are constructed for genes and Robinson-Foulds distances are calculated between gene trees. A graph is constructed with genes as nodes and edge weights from the distances. The strongest edges are then tested for combinability and combined if possible. The final nodes in the graph are the final clusters (i.e., clusters that cannot be combined further).

In a likelihood framework, information criteria are commonly used to accommodate and penalize for the increase in the number of parameters to prevent overfitting. The Akaike Information Criterion (AIC; Akaike 1973), the AIC with the correction for small dataset size (AICc; see Burnham and Anderson 2003), and the Bayesian Information Criterion (BIC; Schwarz 1978) may all be used to compare likelihood scores that are produced from different numbers of parameters. Each of these criteria has different assumptions and different potential utility. Here, we examine the differences in considering AICc and BIC. For both information criteria we use the total number of aligned sites as *n*.

The number of parameters for a single gene in a phylogenetic analysis include those for the molecular model (e.g., *GTR =* 8, 5 for substitution rates and 3 for stationary nucleotide frequencies, with an additional 1 when including gamma-distributed rate variation), and the branch lengths of the unrooted phylogenetic tree (2*n* − 3). There are several ways by which multiple genes may be combined. For example, often molecular models can vary between these genes (or partitions of genes). It is possible to test whether the genes should share models and programs exist to conduct such tests (e.g. PartitionFinder: Lanfear et al. 2016). If each gene region has its own molecular model, then for a multi-gene dataset the number of parameters contributed by molecular models would be the sum of the number of molecular model parameters for each gene. However, for a concatenated analysis of *x* genes, the number of parameters contributed by the branch lengths varies depending on their treatment: shared (2*n* − 3), exactly proportional (‘scaled’ or ‘linked’ (2*n* – 3) + (*x* – 1)), and independent (‘unlinked’; (2*n* – 3) × *x*). Here, we considered the molecular models to be independent between gene regions and tested both scaled and independent branch lengths, as these two methods accommodate rate variation among genes.

With these considerations, the tree comparison calculation proceeded with the following steps. For each gene of interest, calculate the information criterion statistic of the ML gene tree. Next, sum the information criterion statistic for the set of genes being tested. Further, concatenate the genes and calculate the information criterion for the ML tree. The genes may have different model parameters or branch lengths (shared, scaled, or independent), but they share the same topology. Lastly, compare the values of the information criterion for the summed gene trees and the concatenated genes. If the concatenated genes have a lower value of the information criterion than the summed gene trees, accept the combined genes and continue to the next comparison. If genes are already a member of a merged set, then compare the new gene to the merged set. Given this procedure, our algorithm is a greedy clustering method. These methods are implemented in an open source python package, phyckle, available at https://github.com/FePhyFoFum/phyckle.

### Simulations

We verified the performance of the combinatorial method using a variety of simulations across tree depths, branch length heterogeneity, topological variation, and model variation. Each simulation is described below. In general, we attempted to simplify the simulations in order to isolate the specific element being tested in order to better describe the expected behavior. While alignments were simulated under differing models, all clustering tests were conducted using GTR+G as this is typical of empirical analyses. For all simulations below, trees were simulated using **`pxbdsim`** from the phyx package (Brown, Walker, and Smith 2017) and alignments were generated using INDELible (Fletcher and Yang 2009).

#### Comparing information criteria and branch length models

In order to determine the efficacy of different information criteria as well as different branch length models, we conducted several simulation analyses. For each simulation, we generated a tree from a pure birth model with 25 tips and then three gene regions under JC model of evolution with 1000 sites each. This analysis was conducted with 100 replicates. While the JC model of evolution is, perhaps, overly simplistic, we aimed to isolate the factors that caused genes to be considered separate or combined. We test more complex models below. Tree heights were tested for 0.05, 0.25, 0.75, and 1.25. We also conducted tests where branches could vary between gene regions. For each gene region, the species tree branch lengths were perturbed randomly with a sliding window *w* of 0.01, 0.05, and 0.1, so *U* (*x* − *w, x* + *w*). We examined scaled and independent branch length models with both BIC and AICc.

#### Examining the impact of branch differences

To test the impact of having different underlying topologies between gene regions, we simulated a pure birth tree of 25 tips and a tree depth of 0.5 and simulated two gene regions under this model. Then for one additional gene region, we chose one node randomly and swapped nearest neighbors and then simulated gene regions. This resulted in three gene regions with two different underlying topologies. The difference in the underlying topologies varied from one swapping move to five swapping moves. All genes trees also had branch lengths perturbed with a sliding window *w* of 0.01 as described above.

#### Examining the impact of different models on different genes

In order to examine whether different models may cause the gene regions to be considered separate we conducted similar simulations to those described above but with distinct substitution models applied to individual gene regions. Two gene regions were simulated for each of three substitution models (i.e., six gene regions total), each with 1000 bases and the same underlying pure birth topology of 25 taxa and tree depth of 0.5. Branch length heterogeneity varied from 0.01, 0.05, and 0.1 using the sliding window perturbation above. The first two gene regions were evolved under JC, the second set of two gene regions under HKY with *κ* = 2.5, proportion of invariable sites 0.25, *Γ* = 0.5, number of *Γ* categories 10, and state frequencies of 0.2,0.3,0.1,0.4 for A, C, G, and T, respectively, and the third set of two gene region under HKY with *κ* = 1.5, proportion of invariable sites 0.25, *Γ* = 0.5, number of *Γ* categories 10, and state frequencies of 0.1,0.4,0.3,0.2. Two gene regions were simulated for each model in order to verify that those two continued to be clustered together regardless of how the separate models clustered. This test was not intended to be comprehensive as variation in molecular models in relation to information criteria has already been thoroughly explored (e.g., Lanfear et al. 2016; Seo and Thorne 2018). Instead, we aimed to better understand the conditions under which variation in molecular model would result in consideration as separate datasets.

#### Examining the impact of missing taxa

Because genes often do not have completely overlapping taxa, we conducted simulations where some taxa may be missing from each gene region. For these simulations, 25 taxon pure birth trees were generated and three gene regions of 1000 bases each were simulated. Then from one to three tips were randomly removed from one gene (i.e. the aligned sequences of these taxa were replaced with strings of missing data, coded as “-”). We also conducted simulations where from one to three tips were randomly removed from each of the three genes. Random taxa were removed from each gene and so some genes would have the same taxa removed and others would not. All genes trees also had branch lengths perturbed with branch length heterogeneity of 0.01 as described above.

### Empirical Demonstration

For demonstration purposes, we did not conduct exhaustive testing of combinability of the entire Wickett et al. (2014) dataset. Instead, we conducted these tests on the set of 90 genes that supported the magnoliids as sister to the eudicots with greater than 2 lnL versus alternative relationships (Fig. 2). These control methods echo the classes of filtering evoked in Chen, Liang, and Zhang (2015), that of non-specific data filtering (branch length, support values) and ‘node-control’ (outgroup relationships, eudicot relationships).

**Figure 2.**
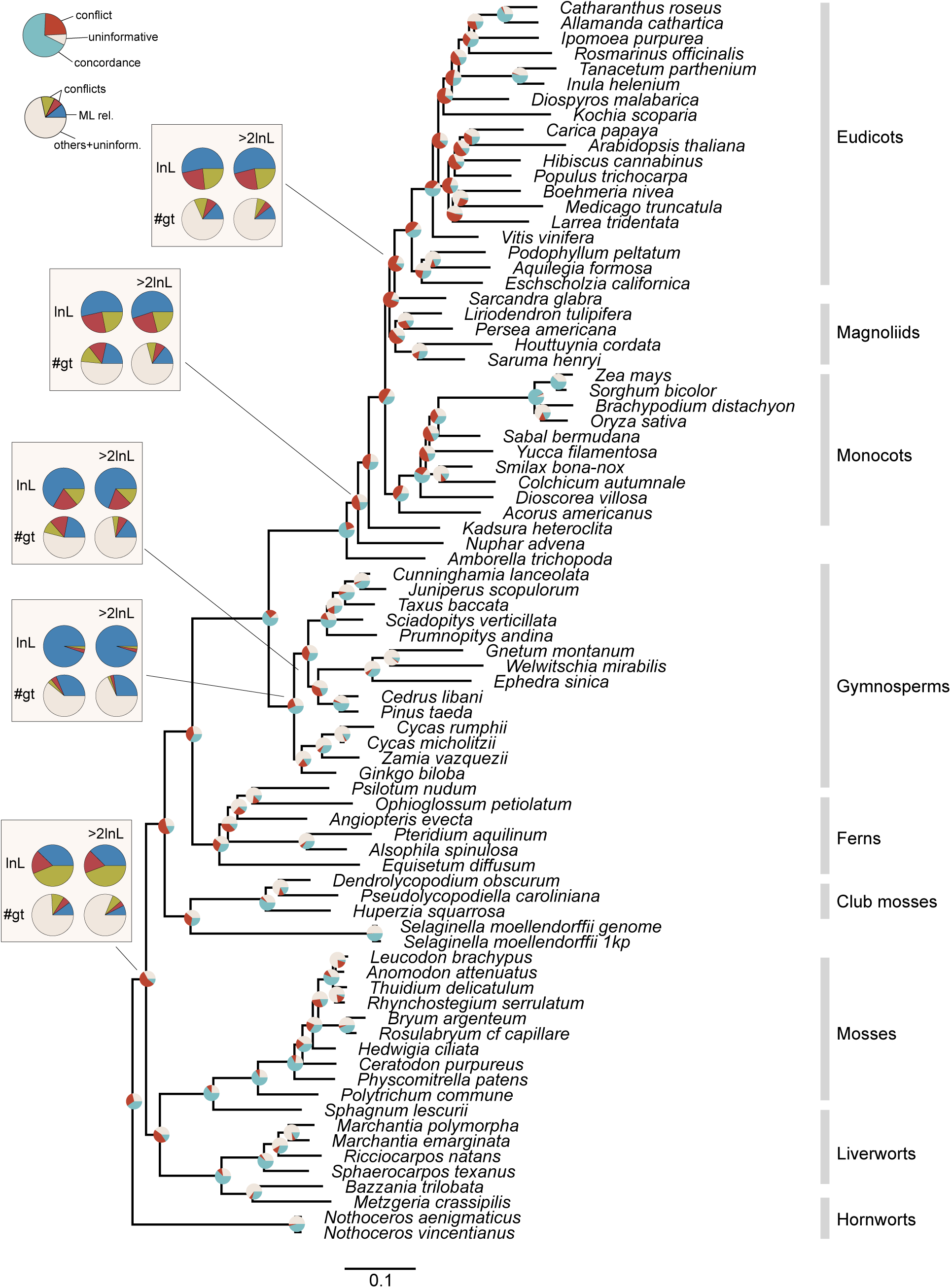
Phylogeny of land plants with pie charts at nodes illustrating conflict, concordance, and informativeness of the gene tree set without any filtering. Inset boxes show summed differences in log likelihoods (top row) and the number of gene trees (bottom row) that support the relationship shown in the tree and the dominant conflicting relationships. Right pie charts in the inset box show results when only differences greater than 2 log likelihoods are considered. See also Table 1.

Clustering analyses were conducted using IQ-TREE with AICc and the **`-spp`** option for scaled branch lengths partitions, as simulations demonstrated that it split the most accurately based on conflicting topologies (see *Results*). We also only considered clustering gene regions that shared at least 70% of their taxa. We refer to our method as the COMBination of datasets (COMB) method. Because our approach bears conceptual similarity to algorithms used to estimate the optimal partitioning schemes (e.g. PartitionFinder, Lanfear et al. 2012, 2016), but additionally considers topological incongruence, we compared combinable subsets to those recommended by the implementation of the PartitionFinder algorithm in IQ-TREE (Kalyaanamoorthy et al. 2017, referred to as MERGE here). Both methods use information criteria (in this case AICc) to ensure direct comparison, in contrast to approaches based on hierarchical LRTs (Leigh et al. 2008). We compared the results of our analyses to the PartitionFinder ‘greedy’ algorithm implemented in IQ-TREE using the option **`-m MERGE`**, specifying the GTR+G model and assessing partitions with the edge-linked proportional model with **`-spp`**. In each case AICc was used for a direct comparison to the results of our method.

## Results

### Conflict analyses

We compared gene trees (Fig. 1) based on the concatenated maximum likelihood (ML) analysis from Wickett et al. (2014) and found that both gene tree conflict and support (Fig. 2) varied through time with support increasing toward the present (Fig. 3). We aimed to resolve contentious relationships, with a focus on those that have either been debated in the literature or been considered important in resolving key evolutionary questions, to the best of the ability of the underlying data (Table 1).

**Figure 3.**
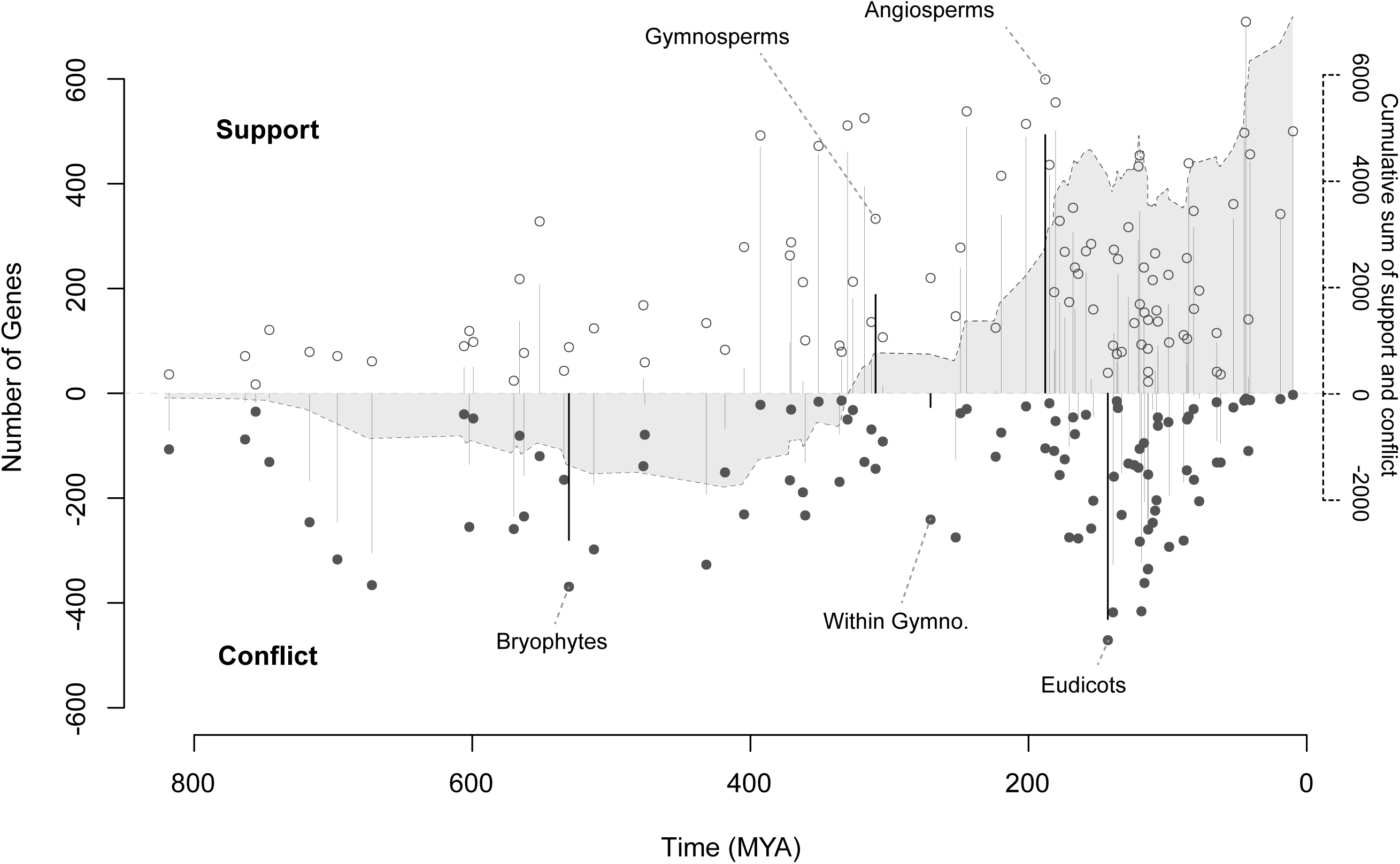
Examination of support and conflict in relation to time across all nodes with node ages taken from TimeTree (Hedges, Dudley, and Kumar 2006; Hedges et al. 2015). The differences between support and conflict are noted with vertical lines. The cumulative sum of support and conflict through time is noted in solid grey. Focal nodes from Fig. 2 are identified.

The massive scale of genomic datasets can cause substantial noise that is often difficult to identify analyzing the entire dataset. When analyzing specific genes, we found that several conflicting relationships were the result of systematic error in the underlying data. In order to minimize the impact of systematic error on the estimation of relationships, we excluded obvious errors where possible. For example, we found 258 of 852 gene trees (30.3%) contained non-land plant taxa that fell within the land plants. While these errors may not impact the estimation of relationships within eudicots, they will impact the estimation of relationships at the origin of land plants. Therefore, we excluded gene trees for which there was not previously well established monophyly of the focal taxa (i.e., involving the relationship of interest). We also identified 68 gene trees that possessed very long estimated branch lengths (> 2.5 expected substitutions per site). We conservatively considered these to contain potential errors in homology (Yang and Smith 2014). While these genes demonstrate patterns associated with systematic error, they also likely contain information for several relationships. However, some error may be the result of misidentified orthology that will mislead estimation of phylogenetic relationships, even if this error may not impact all relationships inferred by the gene. Therefore, to minimize sources of systematic error, we took a conservative approach and excluded these genes from additional analyses.

We explored both numbers of gene trees and differences in log-likelihoods for several key relationships. In some cases, both number of gene trees and differences in log-likelihood support the same resolution, as was the case for the monophyly of Gymnosperms. However, other relationships are more equivocal or contradictory. For example, the relationships among bryophyte lineages are not well supported and equivocal (Table 1).

### Nested analyses

Given the variation in support and conflict through time (Fig. 3), many genes that contain signal for a relationship may disagree with the resolution at other nodes. To examine these patterns of nested conflict, we examined the genes that support the resolution of the eudicot relationships (Fig. 4). In a set of 127 genes which supported the eudicot relationships recovered in the original ML analysis, 90 survived filtering for outgroup placement, branch length, and support with a statistically significant difference in lnL (> 2; Edwards 1984). 57 of these genes displayed the monophyly of gymnosperms, and among those 57 only 23 displayed a sister relationship between pines and Gnetales.

**Figure 4.**
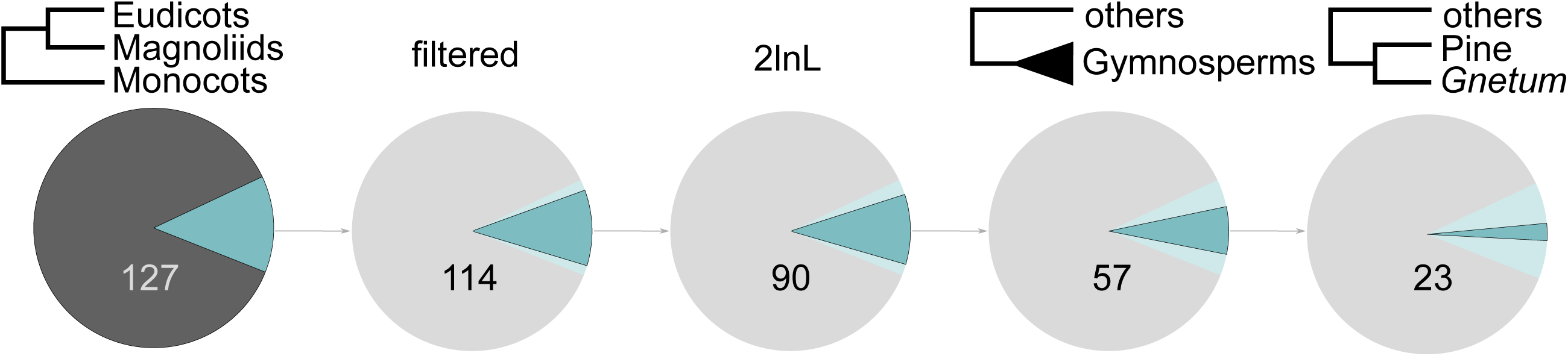
Nested patterns of support with genes associated with the resolution of eudicots. From left to right are shown the genes that support eudicots as sister to magnoliids (far left), those genes filtered as not having any outgroup errors or long branch lengths, those genes that support the resolution by at least 2lnL, those genes that displayed monophyletic gymnosperms, and finally those genes that displayed the Gnepine relationship.

### Simulations of combinability

The combinability procedure described here consists of two components: the information criterion for testing model complexity and the hill-climbing greedy clustering algorithm. First, we conducted analyses to compare the performance of the different information criteria measures (Fig. 5). In our tests, BIC with scaled branch lengths performed the best overall while AICc with scaled branch lengths performed well when branch length heterogeneity was low but poorly when branch length heterogeneity was medium to high. AICc with independent branch lengths tended to overfit (i.e., recover more clusters than truly existed) when tree depths were higher but was more consistent across a range of branch length heterogeneity than any other information criterion. BIC with independent branch lengths (not shown) resulted in underfitting with all genes considered as a single cluster and therefore was not considered further. High branch length heterogeneity generally resulted in overfitting. Because of the propensity of AICc with independent branch lengths to erroneously split clusters with both increasing tree depth and low levels of branch length heterogeneity, we did not consider it further.

**Figure 5.**
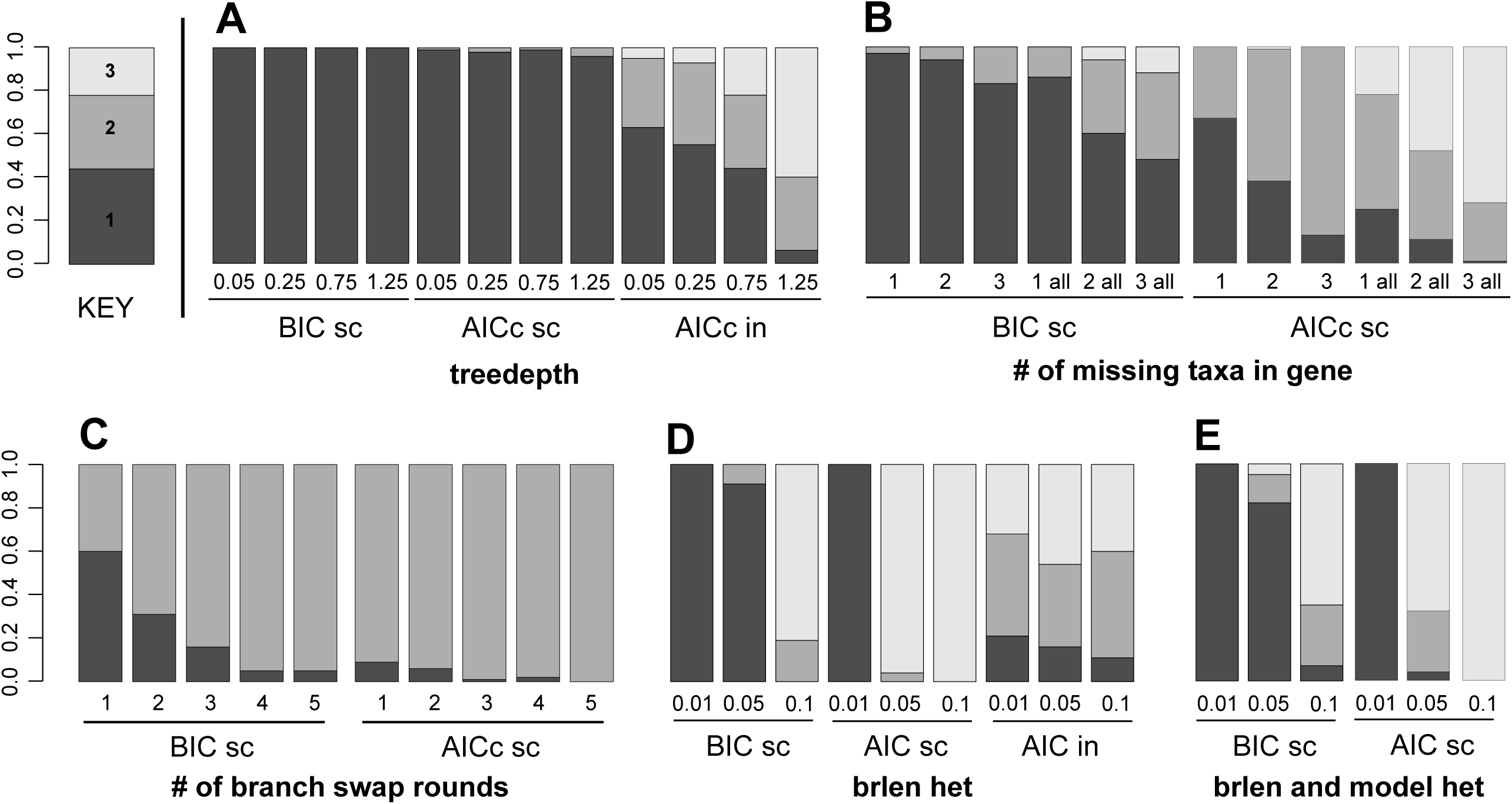
Simulations of clustering behavior for the information criteria-based clustering under different models and data perturbation. ‘sc’ indicates that branch lengths are scaled (proportional) between gene regions, while ‘in’ indicates independent branch lengths. A performance of varied tree depths for three gene regions simulated on the same topology. Ideally all would recover one cluster. B performance for decreasing taxon overlap, with three gene regions simulated on the same topology but with one gene missing 1-3 taxa or with all genes missing 1-3 taxa. Ideally all would recover one cluster. C ability to detect topological differences amongst three gene regions, with two simulated on one topology and one simulated on a topology 1-5 NNI moves away. Ideally all would recover two clusters. D performance of varied branch length heterogeneity for three gene regions simulated on the same topology using the same model. Ideally all would recover one cluster. E performance for three gene regions simulated under different models and increasing branch length heterogeneity on the same topology. Ideally all would recover one cluster.

Phylogenomic datasets often have only partially overlapping taxa sets for each gene, therefore we tested the influence of this in two ways (Fig. 5B). First, we randomly removed from one to three taxa for a single gene. Note that genes otherwise share a topology, and so are best described by a single cluster. The results demonstrate that the procedure tends to overfit as the number of missing taxa increases. AICc with scaled branch lengths was highly sensitive to missing taxa, with between 33% and 87% overfitting for missing taxa in one gene and only one replicate correctly recovering one cluster for the highest number of missing taxa in all genes. BIC with scaled branch lengths was less sensitive to missing taxa, with between 4% and 12% overfitting for missing taxa in one gene, and up to 52% overfitting for missing taxa in all genes. These simulations do not cover instances where there is little to no overlap between datasets. Those scenarios should be explored further.

The results above involve simulations with same underlying species tree topology for each gene. We also examined the ability for the procedure to correctly break up gene regions when underlying topologies differed (Fig. 5C). As the simulations were conducted with two topologies differing from one to five NNIs, we expected the procedure to identify two clusters. We found that AICc with scaled branch lengths was much more sensitive to topological differences, with a highest error of 9% of replicates, and perfect recovery at five NNIs. BIC with scaled branch lengths tended to underfit, with error rates up to 60%, and producing two clusters in 5% of replicates even at five NNIs.

We found that branch length heterogeneity on a common topology is difficult to model (Fig. 5D). AICc with independent branch lengths has the flexibility to model all of the data in a single cluster but overestimates the number of clusters in more than 80% of simulations across all levels of branch length heterogeneity. Scaled branch length models perform better, especially when branch length heterogeneity is low, but are always overfit when heterogeneity is excessive.

While isolating the behavior of the information criteria in relation to tree depth and branch length heterogeneity is helpful, it is likely that most datasets will have variation in substitution models between genes as well (Fig. 5E). We found that the BIC with scaled branch lengths was mostly robust to model variation except in the presence of large branch length heterogeneity (i.e., 10% of total tree height). AICc with scaled branch lengths was prone to overfitting based on model discrepancies, particularly with increasing branch length heterogeneity, correctly recovering one cluster in all replicates with branch length heterogeneity of 0.01, but incorrectly recovering three clusters in all replicates with branch length heterogeneity of 0.01. The discrepancy between the branch length heterogeneity of 0.1 in this analysis and the one above reflects that there were six genes simulated in this case with two for each model versus three gene regions as above.

### Empirical combinability of genes

No method or gene set supported the concatenation of all genes that supported the focal eudicot relationship. The COMB method on the ‘CombinedSet’ supported concatenation of three sets with two genes each and one set with three genes. There was sensitivity to the order of comparisons in the graph as weighted RF had partially overlapping but different results with five sets of two genes each. The phylogenies produced from concatenation did not result in conflict with the monophyly of eudicots and magnoliids but only three of four supported that relationship with greater than 80% support from the SH-aLRT. MERGE with AICc produced 18 clusters, of which one contained four genes, four contained three genes, and 13 contained two genes. We inferred trees from the concatenated sequences of each cluster and assessed their topologies: only 10 supported the monophyly of eudicots and magnoliids with greater than 80% support from the SH-aLRT. MERGE supported partition merging for a much greater number of genes than COMB supported combination.

## Discussion

### Conflict analysis

Several contentious relationships show strong contrast between the number of genes supporting the relationship, the number of genes strongly supporting the relationship (>2 lnL), the ΔlnL supporting the relationship, and the ΔlnL of genes that strongly support the relationship. Our analyses demonstrated that the differences in the number of gene trees supporting relationships and the difference in the summed ΔlnLs can provide insight into the cause for discordance between concatenated ML analyses and coalescent analyses (Table 1).

### Nested analysis

Filtering genes by the specific relationship they display provides an opportunity to examine nested conflicts (i.e., subsets of genes that do not conflict in one relationship may conflict in another). Furthermore, if conflict was reduced as a result of filtering, concatenation may be more tenable on such filtered datasets. However, our nested conflict analyses demonstrated significant conflict and variation in the support for different relationships (Fig. 4) and that filtering genes based on specific relationships did not reduce conflict in other parts of the tree. While filtering genes may provide some means for lessening some systematic errors (Brown and Thomson 2017), or reducing some conflict (the question-specific ‘node-control’ approach of Chen, Liang, and Zhang (2015)) our analyses suggest that it will not likely solve general problems regarding conflicting genes.

### Combinability of genes

It is perhaps naïve to expect a single gene to have high support throughout a large part of the Tree of Life (see Penny et al. (1990); MUTOG: the ‘Myth of a Universal Tree from One Gene’). For this reason, some researchers have argued that concatenating genes effectively combines data informative at various scales and so provides the necessary information to better resolve deep and shallow nodes (e.g., Mirarab, Bayzid, et al. 2014). Despite the potential benefits of concatenation (i.e., amplifying weak phylogenetic signal), the underlying model of evolution for a concatenated analysis assumes topological concordance among gene tree histories. Extensive gene conflicts should often violate these assumptions. Filtering genes could be one means of reducing conflict, though our filtered analyses demonstrated that conflict remained in other parts of the tree. Whether genes should be combined for a concatenated analysis has been discussed at length (Huelsenbeck, Bull, and Cunningham 1996; Leigh et al. 2008; Seo and Thorne 2018; Theobald 2010; Walker, Brown, and Smith 2018) and, in addition to traditional tests, Bayesian methods have recently been developed to address some of these issues (Neupane et al. 2018). However, due to the large scale of genomic datasets, Bayesian methods are often computationally intractable.

We described a simple and greedy method meant to examine whether genes should be combined based on information criteria and validated its performance through simulation. Our approach builds on a long history of tests for the combinability of phylogenetic data (Huelsenbeck, Bull, and Cunningham 1996; Cunningham 1997; Shimodaira and Hasegawa 1999; Ané et al. 2006; Leigh et al. 2008), for the merging of multiple partitions in substitution models (Lanfear et al. 2012, 2016), and for recombination based on detection of multiple tree signals within genes (Kosakovsky Pond et al. 2006a, 2006b; Ané 2011). Like the latter methods, we allow free estimation of gene tree topologies and use information criterion to assess combinability of gene regions. Our approach also bears resemblance to statistical binning approaches with the addition of information theoretic measures but without the intention of species tree analysis of the clusters (Mirarab, Bayzid, et al. 2014).

Simulations demonstrated that our approach performed well with clustering success decreasing with increasing tree depth and increasing branch length heterogeneity (Fig. 5). Simply put, trees that were more different were less likely to be clustered together. Based on these results, we find that our method provides a feasible, if conservative (e.g., tending to split), approach to examine combinable subsets of genes. Despite the shortcuts employed, however, it involves long computational times and is intractable for large datasets.

Using this heuristic, we tested combinability of the subset of the genes from Wickett et al. (2014) that supported magnoliids sister to eudicots as inferred in the original ML analysis. In general, only a very small set of genes supported concatenation. Given the extensive underlying gene tree conflict that remains even after filtering for a focal relationship (Fig. 4) this should be expected as simulations demonstrated that our approach using AICc with scaled branch lengths is very sensitive to topological heterogeneity.

Because of the extensive gene tree conflict within datasets and the improbable nature of supporting combining genes that differ extensively in topology, we caution the use of the approach outlined here for broad and unsupervised application. We argue that more targeted, data-centric approaches to large phylogenomic datasets are more illuminating regarding the support for specific relationships than broad concatenation. Other considerations are also important such as the impact of missing data, the order of comparisons, and the impact of different information criteria and branch length models. For example, simulations demonstrated sensitivity to missing taxa while the empirical tests suggest a clustering of genes when there was very poor overlap. Concatenation is a common approach for analyzing phylogenomic analyses and while concatenation can be helpful for exploratory inference to identify dominant signal, it is not capable of addressing specific and contentious relationships.

### Implications for plant phylogenetics

The results presented here provide strong support for several relationships that have long been considered contentious and indicate probable resolutions for others. For example, we found support for *Amborella* being sister to the rest of angiosperms and that gymnosperms are monophyletic. Several relationships (e.g., the hornworts, liverworts, and mosses) lack enough information to confidently accept any of the alternative resolutions. Rather than being dismayed at this apparent failure, we regard this lack of signal as extremely valuable information, as it informs where future effort should be focused. Though we identified the relationship that was more strongly supported by the data (Table 1), the differences between the alternatives were so slight that the current dataset is likely unable to confidently resolve this debate and conducting additional analyses with expanded taxa and gene regions is warranted.

Among the strongly supported hypotheses, the placement of *Amborella* continues to be a point of major contention within the plant community. *Amborella* is a tropical tree with relatively small flowers, while the Nymphaeales are aquatic plants with relatively large flowers. The resolution of these taxa in relation to the remainder of the flowering plants will inform the life history of early angiosperms (Feild et al. 2004) as well as the lability of life history and floral traits. Our results suggest *Amborella* is sister to all other extant angiosperms and imply that rates of evolution need not be particularly fast in order to understand the morphological differences between a tropical tree (*Amborella*) and water lilies (Nymphaeales). Strong support for the monophyly of gymnosperms implies that the morphological disparity of extant gymnosperm taxa, including the especially diverse Gnetales, emerged post-divergence from the angiosperm lineage. This reinforces analyses of LEAFY homologs, which recover gymnosperm paralogs as monophyletic groups (Sayou et al. 2014), and also lends support to shared characteristics between Gnetales and angiosperms resulting from convergent evolution (Bowe, Coat, and dePamphilis 2000; Hansen et al. 1999).

For contentious relationships only weakly supported here, there are several biological questions that will be answered once/if these are confidently resolved. The data and analyses presented here suggest a difficulty to distinguish between monophyletic bryophytes and hornworts as sister to all other land plants. Recovering hornworts as sister is consistent with some studies (Nickrent et al. 2000; Nishiyama and Kato 1999) but contradicts the results of others (Cox et al. 2014; Karol et al. 2010; Qiu et al. 2006), including some but not all results of a recent re-analysis of this dataset (Puttick et al. 2018). If this position were supported with additional data, it implies that the absence of stomata in liverworts and some mosses is a derived state resulting from loss of the trait, suggests a single loss of pyrenoids in non-hornwort land plants (but see Villarreal and Renner 2012), and questions some inferences on putative morphological synapomorphies between hornworts and vascular plants, such as a nutritionally-independent sporophyte (Qiu et al. 2006; Wickett et al. 2014). However, more data are required to determine the relative support between these two dominant hypotheses. Among gymnosperms, these data suggest that Gnetales are sister to pines (the “Gnepine” hypothesis; Chaw et al. 2000), further supporting the lability and rapid evolution of morphological disparity within the group. Finally, magnoliids are inferred as sister to the eudicot lineages, which has implications on the origin and divergence times of eudicots and monocots.

Despite the ability of the methods explored here to accommodate the underlying gene tree uncertainty, our results depend on the information available in the underlying dataset. While this dataset is not comprehensive, it *does* represent extensive sequencing of transcriptomes and genomes for the taxa included. We can say, with confidence, what these data support or do not support, but different datasets (e.g., based on different taxa, different homology analyses) may have stronger signal for relationships that are resolved more equivocally here. We recommend analyzing these future datasets with an eye toward hypotheses of specific phylogenetic relationships. Our novel approach provides insight into several of the most contentious relationships across land plants and is broadly applicable among different groups. Approaches that ascertain the support for alternative resolutions should be used to resolve contentious branches across the Tree of Life.

### Concatenation, coalescent, and a data-centric middle way

A panacea does not currently exist for phylogenomic analyses. Some researchers aim to determine the relative support for contentious relationships. Others want to construct a reasonable, if not ideal, phylogeny for downstream analyses. Others still may be primarily interested in gene trees. Here, we suggest that more detailed and data-centric analyses of the gene trees and specific relationships will yield more useful results concerning the resolution of relationships.

Our results also speak to several common analyses conducted on phylogenomic datasets. The underlying conflict identified by many researchers (Wickett et al. 2014; Puttick et al. 2018) suggests that concatenation, while helpful for identifying the dominant signal, should not be used to address contentious nodes. Our targeted exploration of the combinability of gene regions found that very few genes are optimally modelled by concatenation, even when filtering on those genes that support a relationship. However, our analyses of combinability leave many unanswered questions. For example, how should we adequately address the problem of low signal when gene tree conflict is high, and concatenation is statistically broadly unsupported? Some have suggested binning genes based on similarity to amplify signal and feeding the clusters into a coalescent analysis (Mirarab, Bayzid, et al. 2014). This may or may not violate assumptions of the coalescent (e.g., are concatenated genes linked or unlinked) but even the assumptions are not violated, these analyses do not facilitate better understanding of underlying conflict within the dataset.

The most common alternatives to concatenated supermatrix analyses are coalescent species tree approaches, which often accommodate one major source of conflict (usually ILS) across gene trees (Mirarab, Reaz, et al. 2014). The most sophisticated model-based coalescent approaches are often not computationally tractable for phylogenomic analyses because of the large sizes of the datasets (Ané et al. 2006; Boussau et al. 2013). As a result, most phylogenomic analyses that accommodate ILS use quartet methods (e.g., ASTRAL) that, while fast and effective, do not account for multiple sources of conflict and make several other assumptions that may or may not be reasonable given the dataset (e.g. equal weighting of gene trees). Some researchers have suggested filtering the data to include only those genes that conflict due to ILS (Knowles et al. 2018; Huang et al. 2017) or that agree with accepted relationships or specific relationships to be tested (Chen, Liang, and Zhang 2015; Doyle et al. 2015; Smith, Brown, and Walker 2018). However, for datasets with a broad scope, several processes may be at play throughout the phylogeny and it may not be possible to filter based on a single underlying process. Furthermore, concatenation analyses alone do not provide information regarding the nature of underlying conflict within the dataset or the relative informativeness or lack thereof for genes across a dataset.

While a single species tree may be necessary for some downstream analyses, they obfuscate the biological realities that underlie these data. Even if both broad concatenation and coalescent analyses produced accurate trees for a specific dataset, more detailed and data-centric analyses would provide important details regarding the relative support and conflict underlying these rich genomic datasets. By uncovering the support and lack thereof, we can determine the limits of our data, identify troublesome phylogenetic relationships that require more attention, and put to rest debates over specific relationships (at least regarding specific datasets). The approach we adopt here is akin to the ‘hypothesis-control’ method of Chen, Liang, and Zhang (2015), but instead of relying on the results of typical inference on the filtered subsets, we profile the signal for different resolutions and processes within them. Though additional work is necessary to translate these results to species tree analyses, these approaches along with others over the last few years provide first steps (e.g., Shen, Hittinger, and Rokas 2017; Brown and Thomson 2017; Walker, Brown, and Smith 2018). Over the last decade, massive data sets have been collected and analyzed and overwhelmingly researchers have identified widespread conflict and heterogeneity. We argue that our analyses should reflect these discoveries and that by pursuing, in more detail, the underlying signal within these rich datasets, we will better understand the patterns and processes that shape the tree of life.

## Acknowledgments

This work was supported by funding from NSF DEB 1354048 (J.F.W. and S.A.S.), NSF AVATOL 1207915 (J.W.B. and S.A.S.) and the Woolf Fisher Trust (N.W.-H.). We would like to thank the AE and reviewers for providing thoughtful and helpful comments. We also appreciate comments from Ning Wang, Caroline Parins-Fukuchi, Diego Alvarado Serrano, Greg Stull, Drew Larson, Hector Fox, and Richie Hodel.

## Literature Cited

Ahrenfeldt, Johanne, Carina Skaarup, Henrik Hasman, Anders Gorm Pedersen, Frank Møller Aarestrup, and Ole Lund. 2017. “Bacterial Whole Genome-Based Phylogeny: Construction of a New Benchmarking Dataset and Assessment of Some Existing Methods.” BMC Genomics 18 (1). BioMed Central: 19.

Akaike, Hirotogu. 1973. “Information Theory and an Extension of the Maximum Likelihood Principle.” In Second International Symposium on Information Theory, edited by Petrov P. N. and Csaki F., 267–81. Akademiai Kiado.

Ané, Cécile. 2011. “Detecting Phylogenetic Breakpoints and Discordance from Genome-Wide Alignments for Species Tree Reconstruction.” Genome Biology and Evolution 3 (February): 246–58. https://doi.org/10.1093/gbe/evr013.

Ané, Cécile, Bret Larget, David A Baum, Stacey D Smith, and Antonis Rokas. 2006. “Bayesian Estimation of Concordance Among Gene Trees.” Molecular Biology and Evolution 24 (2). Oxford University Press: 412–26.

Boussau, Bastien, Gergely J Szöllősi, Laurent Duret, Manolo Gouy, Eric Tannier, and Vincent Daubin. 2013. “Genome-Scale Coestimation of Species and Gene Trees.” Genome Research 23 (2). Cold Spring Harbor Lab: 323–30.

Bowe, L Michelle, Gwénaële Coat, and Claude W. dePamphilis. 2000. “Phylogeny of Seed Plants Based on All Three Genomic Compartments: Extant Gymnosperms Are Monophyletic and Gnetales’ Closest Relatives Are Conifers.” Proceedings of the National Academy of Sciences 97 (8). National Acad Sciences: 4092–7.

Brown, Jeremy M, and Robert C Thomson. 2017. “Bayes Factors Unmask Highly Variable Information Content, Bias, and Extreme Influence in Phylogenomic Analyses.” Systematic Biology 66 (4). Oxford University Press: 517–30.

Brown, Joseph W, Joseph F Walker, and Stephen A Smith. 2017. “Phyx: Phylogenetic Tools for Unix.” Bioinformatics 33 (12). Oxford University Press: 1886–8.

Burnham, Kenneth P, and David R Anderson. 2003. Model Selection and Multimodel Inference: A Practical Information-Theoretic Approach. Springer Science & Business Media.

Chaw, Shu-Miaw, Christopher L Parkinson, Yuchang Cheng, Thomas M Vincent, and Jeffrey D Palmer. 2000. “Seed Plant Phylogeny Inferred from All Three Plant Genomes: Monophyly of Extant Gymnosperms and Origin of Gnetales from Conifers.” Proceedings of the National Academy of Sciences 97 (8). National Acad Sciences: 4086–91.

Chen, Meng-Yun, Dan Liang, and Peng Zhang. 2015. “Selecting Question-Specific Genes to Reduce Incongruence in Phylogenomics: A Case Study of Jawed Vertebrate Backbone Phylogeny.” Systematic Biology 64 (6). Oxford University Press: 1104–20.

Cox, Cymon J., Blaise Li, Peter G. Foster, T. Martin Embley, and Peter Civán. 2014. “Conflicting Phylogenies for Early Land Plants Are Caused by Composition Biases Among Synonymous Substitutions.” Systematic Biology 63 (2). Oxford University Press: 272–79. http://dx.doi.org/10.1093/sysbio/syt109.

Cunningham, C W. 1997. “Can three incongruence tests predict when data should be combined?” Molecular Biology and Evolution 14 (7): 733–40. https://doi.org/10.1093/oxfordjournals.molbev.a025813.

Doyle, Vinson P, Randee E Young, Gavin JP Naylor, and Jeremy M Brown. 2015. “Can We Identify Genes with Increased Phylogenetic Reliability?” Systematic Biology 64 (5). Oxford University Press: 824–37.

Dunn, Casey W, Andreas Hejnol, David Q Matus, Kevin Pang, William E Browne, Stephen A Smith, Elaine Seaver, et al. 2008. “Broad Phylogenomic Sampling Improves Resolution of the Animal Tree of Life.” Nature 452 (7188). Nature Publishing Group: 745.

Edwards, Anthony William Fairbank. 1984. Likelihood. CUP Archive.

Edwards, Scott V. 2009. “Is a New and General Theory of Molecular Systematics Emerging?” Evolution 63 (1). Wiley Online Library: 1–19.

Edwards, Scott V, Liang Liu, and Dennis K Pearl. 2007. “High-Resolution Species Trees Without Concatenation.” Proceedings of the National Academy of Sciences 104 (14). National Acad Sciences: 5936–41.

Edwards, Scott V, Zhenxiang Xi, Axel Janke, Brant C Faircloth, John E McCormack, Travis C Glenn, Bojian Zhong, et al. 2016. “Implementing and Testing the Multispecies Coalescent Model: A Valuable Paradigm for Phylogenomics.” Molecular Phylogenetics and Evolution 94. Elsevier: 447–62.

Feild, Taylor S, Nan Crystal Arens, James A Doyle, Todd E Dawson, and Michael J Donoghue. 2004. “Dark and Disturbed: A New Image of Early Angiosperm Ecology.” Paleobiology 30 (1). BioOne: 82–107.

Feuda, Roberto, Martin Dohrmann, Walker Pett, Hervé Philippe, Omar Rota-Stabelli, Nicolas Lartillot, Gert Wörheide, and Davide Pisani. 2017. “Improved Modeling of Compositional Heterogeneity Supports Sponges as Sister to All Other Animals.” Current Biology 27 (24). Elsevier: 3864–70.

Fletcher, William, and Ziheng Yang. 2009. “INDELible: A Flexible Simulator of Biological Sequence Evolution.” Molecular Biology and Evolution 26 (8). SMBE: 1879–88.

Guindon, Stéphane, Jean-François Dufayard, Vincent Lefort, Maria Anisimova, Wim Hordijk, and Olivier Gascuel. 2010. “New Algorithms and Methods to Estimate Maximum-Likelihood Phylogenies: Assessing the Performance of Phyml 3.0.” Systematic Biology 59 (3). Oxford University Press: 307–21.

Hansen, Andrea, Sabine Hansmann, Tahir Samigullin, Andrey Antonov, and William Martin. 1999. “Gnetum and the Angiosperms: Molecular Evidence That Their Shared Morphological Characters Are Convergent, Rather Than Homologous.” Molecular Biology and Evolution 16 (7). Oxford University Press: 1006–6.

Hedges, S Blair, Joel Dudley, and Sudhir Kumar. 2006. “TimeTree: A Public Knowledge-Base of Divergence Times Among Organisms.” Bioinformatics 22 (23). Oxford University Press: 2971–2.

Hedges, S. Blair, Julie Marin, Michael Suleski, Madeline Paymer, and Sudhir Kumar. 2015. “Tree of Life Reveals Clock-Like Speciation and Diversification.” Molecular Biology and Evolution 32 (4): 835–45. https://doi.org/10.1093/molbev/msv037.

Huang, Huateng, Jeet Sukumaran, Stephen A Smith, and LLacey Knowles. 2017. “Cause of Gene Tree Discord? Distinguishing Incomplete Lineage Sorting and Lateral Gene Transfer in Phylogenetics.” PeerJ PrePrints 5. PeerJ, Inc.: e3489v1.

Huelsenbeck, John P, JJ Bull, and Clifford W Cunningham. 1996. “Combining Data in Phylogenetic Analysis.” Trends in Ecology & Evolution 11 (4). Elsevier: 152–58.

Jarvis, Erich D, Siavash Mirarab, Andre J Aberer, Bo Li, Peter Houde, Cai Li, Simon YW Ho, et al. 2014. “Whole-Genome Analyses Resolve Early Branches in the Tree of Life of Modern Birds.” Science 346 (6215). American Association for the Advancement of Science: 1320–31.

Kalyaanamoorthy, Subha, Bui Quang Minh, Thomas KF Wong, Arndt von Haeseler, and Lars S Jermiin. 2017. “ModelFinder: Fast Model Selection for Accurate Phylogenetic Estimates.” Nature Methods 14 (6). Nature Publishing Group: 587.

Karol, Kenneth G., Kathiravetpillai Arumuganathan, Jeffrey L. Boore, Aaron M. Duffy, Karin DE Everett, John D. Hall, S. Kellon Hansen, et al. 2010. “Complete Plastome Sequences of Equisetum Arvense and Isoetes Flaccida: Implications for Phylogeny and Plastid Genome Evolution of Early Land Plant Lineages.” BMC Evolutionary Biology 10 (1): 321. https://doi.org/10.1186/1471-2148-10-321.

Kluge, Arnold G. 1989. “A Concern for Evidence and a Phylogenetic Hypothesis of Relationships Among Epicrates (Boidae, Serpentes).” Systematic Biology 38 (1). Society of Systematic Zoology: 7–25.

Knowles, L Lacey, Huateng Huang, Jeet Sukumaran, and Stephen A Smith. 2018. “A Matter of Phylogenetic Scale: Distinguishing Incomplete Lineage Sorting from Lateral Gene Transfer as the Cause of Gene Tree Discord in Recent Versus Deep Diversification Histories.” American Journal of Botany 105 (3). Wiley Online Library: 376–84.

Kosakovsky Pond, Sergei L, David Posada, Michael B Gravenor, Christopher H Woelk, and Simon DW Frost. 2006a. “Automated Phylogenetic Detection of Recombination Using a Genetic Algorithm.” Molecular Biology and Evolution 23 (10). Oxford University Press: 1891–1901.

Kosakovsky Pond, Sergei L. 2006b. “GARD: A Genetic Algorithm for Recombination Detection.“ Bioinformatics 22 (24). Oxford University Press: 3096–8.

Lanfear, Robert, Brett Calcott, Simon YW Ho, and Stephane Guindon. 2012. “PartitionFinder: Combined Selection of Partitioning Schemes and Substitution Models for Phylogenetic Analyses.” Molecular Biology and Evolution 29 (6). Oxford University Press: 1695–1701.

Lanfear, Robert, Paul B Frandsen, April M Wright, Tereza Senfeld, and Brett Calcott. 2016. “PartitionFinder 2: New Methods for Selecting Partitioned Models of Evolution for Molecular and Morphological Phylogenetic Analyses.” Molecular Biology and Evolution 34 (3). Oxford University Press: 772–73.

Leigh, Jessica W, Edward Susko, Manuela Baumgartner, and Andrew J Roger. 2008. “Testing Congruence in Phylogenomic Analysis.” Systematic Biology 57 (1). Taylor & Francis: 104–15.

Liu, Liang, Lili Yu, Dennis K Pearl, and Scott V Edwards. 2009. “Estimating Species Phylogenies Using Coalescence Times Among Sequences.” Systematic Biology 58 (5). Oxford University Press: 468–77.

Maddison, Wayne P. 1997. “Gene Trees in Species Trees.” Systematic Biology 46 (3). Oxford University Press: 523–36.

Mirarab, Siavash, Md Shamsuzzoha Bayzid, Bastien Boussau, and Tandy Warnow. 2014. “Statistical Binning Enables an Accurate Coalescent-Based Estimation of the Avian Tree.” Science 346 (6215). American Association for the Advancement of Science: 1250463.

Mirarab, Siavash, Rezwana Reaz, Md S Bayzid, Théo Zimmermann, M Shel Swenson, and Tandy Warnow. 2014. “ASTRAL: Genome-Scale Coalescent-Based Species Tree Estimation.” Bioinformatics 30 (17). Oxford University Press: i541–i548.

Neupane, Suman, Karolina Fucikova, Louise A Lewis, Lynn Kuo, Ming-Hui Chen, and Paul Lewis. 2018. “Assessing Combinability of Phylogenomic Data Using Bayes Factors.” bioRxiv. Cold Spring Harbor Laboratory, 250969.

Nguyen, Lam-Tung, Heiko A Schmidt, Arndt von Haeseler, and Bui Quang Minh. 2015. “IQ-Tree: A Fast and Effective Stochastic Algorithm for Estimating Maximum-Likelihood Phylogenies.” Molecular Biology and Evolution 32 (1). Oxford University Press: 268–74.

Nickrent, Daniel L., Christopher L. Parkinson, Jeffrey D. Palmer, and R. Joel Duff. 2000. “Multigene Phylogeny of Land Plants with Special Reference to Bryophytes and the Earliest Land Plants.” Molecular Biology and Evolution 17 (12): 1885–95. https://doi.org/10.1093/oxfordjournals.molbev.a026290.

Nishiyama, T, and M Kato. 1999. “Molecular Phylogenetic Analysis Among Bryophytes and Tracheophytes Based on Combined Data of Plastid Coded Genes and the 18S rRNA Gene.” Molecular Biology and Evolution 16 (8): 1027–36. https://doi.org/10.1093/oxfordjournals.molbev.a026192.

Penny, D, MD Hendy, EA Zimmer, and RK Hamby. 1990. “Trees from Sequences: Panacea or Pandora’s Box?” Australian Systematic Botany 3 (1). CSIRO: 21–38.

Puttick, Mark N, Jennifer L Morris, Tom A Williams, Cymon J Cox, Dianne Edwards, Paul Kenrick, Silvia Pressel, et al. 2018. “The Interrelationships of Land Plants and the Nature of the Ancestral Embryophyte.” Current Biology 28 (5). Elsevier: 733–45.

Qiu, Yin-Long, Libo Li, Bin Wang, Zhiduan Chen, Volker Knoop, Milena Groth-Malonek, Olena Dombrovska, et al. 2006. “The Deepest Divergences in Land Plants Inferred from Phylogenomic Evidence.” Proceedings of the National Academy of Sciences 103 (42). National Academy of Sciences: 15511–6. https://doi.org/10.1073/pnas.0603335103.

Robinson, David F, and Leslie R Foulds. 1981. “Comparison of Phylogenetic Trees.” Mathematical Biosciences 53 (1-2). Elsevier: 131–47.

Sauquet, Hervé, Maria von Balthazar, Susana Magallón, James A Doyle, Peter K Endress, Emily J Bailes, Erica Barroso de Morais, et al. 2017. “The Ancestral Flower of Angiosperms and Its Early Diversification.” Nature Communications 8. Nature Publishing Group: 16047.

Sayou, Camille, Marie Monniaux, Max H Nanao, Edwige Moyroud, Samuel F Brockington, Emmanuel Thévenon, Hicham Chahtane, et al. 2014. “A Promiscuous Intermediate Underlies the Evolution of Leafy Dna Binding Specificity.” Science 343 (6171). American Association for the Advancement of Science: 645–48.

Schwarz, Gideon. 1978. “Estimating the Dimension of a Model.” The Annals of Statistics 6 (2). Institute of Mathematical Statistics: 461–64.

Seo, Tae-Kun, and Jeffrey L Thorne. 2018. “Information Criteria for Comparing Partition Schemes.” Systematic Biology 67 (4): 616–32. https://doi.org/10.1093/sysbio/syx097.

Shen, Xing-Xing, Chris Todd Hittinger, and Antonis Rokas. 2017. “Contentious Relationships in Phylogenomic Studies Can Be Driven by a Handful of Genes.” Nature Ecology & Evolution 1 (5). Nature Publishing Group: 0126.

Shen, Xing-Xing, Xiaofan Zhou, Jacek Kominek, Cletus P Kurtzman, Chris Todd Hittinger, and Antonis Rokas. 2016. “Reconstructing the Backbone of the Saccharomycotina Yeast Phylogeny Using Genome-Scale Data.” G3: Genes, Genomes, Genetics 6 (12). G3: Genes, Genomes, Genetics: 3927–39.

Shimodaira, Hidetoshi, and Masami Hasegawa. 1999. “Multiple Comparisons of Log-Likelihoods with Applications to Phylogenetic Inference.” Molecular Biology and Evolution 16 (8). Oxford University Press: 1114–4.

Simion, Paul, Hervé Philippe, Denis Baurain, Muriel Jager, Daniel J. Richter, Arnaud Di Franco, Béatrice Roure, et al. 2017. “A Large and Consistent Phylogenomic Dataset Supports Sponges as the Sister Group to All Other Animals.” Current Biology 27 (7):958–67. https://doi.org/https://doi.org/10.1016/j.cub.2017.02.031.

Smith, Stephen A, Joseph W Brown, and Joseph F Walker. 2018. “So Many Genes, so Little Time: A Practical Approach to Divergence-Time Estimation in the Genomic Era.” PloS One 13 (5). Public Library of Science: e0197433.

Smith, Stephen A, Michael J Moore, Joseph W Brown, and Ya Yang. 2015. “Analysis of Phylogenomic Datasets Reveals Conflict, Concordance, and Gene Duplications with Examples from Animals and Plants.” BMC Evolutionary Biology 15 (1). BioMed Central: 150.

Solís-Lemus, Cécile, Claudia and Ané. 2016. “Inferring Phylogenetic Networks with Maximum Pseudolikelihood Under Incomplete Lineage Sorting.” PLOS Genetics 12 (3). Public Library of Science: 1–21. https://doi.org/10.1371/journal.pgen.1005896.

Springer, Mark S, and John Gatesy. 2016. “The Gene Tree Delusion.” Molecular Phylogenetics and Evolution 94. Elsevier: 1–33.

Stamatakis, Alexandros. 2014. “RAxML version 8: a tool for phylogenetic analysis and post-analysis of large phylogenies.” Bioinformatics 30. 1312–1313.

Theobald, Douglas L. 2010. “A Formal Test of the Theory of Universal Common Ancestry.” Nature 465 (7295). Nature Publishing Group: 219.

Villarreal, Juan Carlos, and Susanne S Renner. 2012. “Hornwort Pyrenoids, Carbon-Concentrating Structures, Evolved and Were Lost at Least Five Times During the Last 100 Million Years.” Proceedings of the National Academy of Sciences 109 (46). National Acad Sciences: 18873–8.

Walker, Joseph F, Joseph W Brown, and Stephen A Smith. 2018. “Analyzing Contentious Relationships and Outlier Genes in Phylogenomics.” Systematic Biology syy043. Oxford University Press. https://doi.org/10.1093/sysbio/syy043.

Walker, Joseph F, Ya Yang, Tao Feng, Alfonso Timoneda, Jessica Mikenas, Vera Hutchison, Caroline Edwards, et al. 2018. “From Cacti to Carnivores: Improved Phylotranscriptomic Sampling and Hierarchical Homology Inference Provide Further Insight into the Evolution of Caryophyllales.” American Journal of Botany 105 (3). Wiley Online Library: 446–62.

Walker, Joseph F, Ya Yang, Michael J Moore, Jessica Mikenas, Alfonso Timoneda, Samuel F Brockington, and Stephen A Smith. 2017. “Widespread Paleopolyploidy, Gene Tree Conflict, and Recalcitrant Relationships Among the Carnivorous Caryophyllales.” American Journal of Botany 104 (6). Botanical Soc America: 858–67.

Wen, Dingqiao, Yun Yu, Jiafan Zhu, and Luay Nakhleh. 2018. “Inferring Phylogenetic Networks Using Phylonet.” Systematic Biology 67 (4): 735–40. https://doi.org/10.1093/sysbio/syy015.

Whelan, Nathan V, Kevin M Kocot, Tatiana P Moroz, Krishanu Mukherjee, Peter Williams, Gustav Paulay, Leonid L Moroz, and Kenneth M Halanych. 2017. “Ctenophore Relationships and Their Placement as the Sister Group to All Other Animals.” Nature Ecology & Evolution 1 (11). Nature Publishing Group: 1737.

Wickett, Norman J, Siavash Mirarab, Nam Nguyen, Tandy Warnow, Eric Carpenter, Naim Matasci, Saravanaraj Ayyampalayam, et al. 2014. “Phylotranscriptomic Analysis of the Origin and Early Diversification of Land Plants.” Proceedings of the National Academy of Sciences 111 (45). National Acad Sciences: E4859–E4868.

Xi, Zhenxiang, Liang Liu, Joshua S Rest, and Charles C Davis. 2014. “Coalescent Versus Concatenation Methods and the Placement of Amborella as Sister to Water Lilies.” Systematic Biology 63 (6). Oxford University Press: 919–32.

Yang, Ya, and Stephen A Smith. 2014. “Orthology Inference in Nonmodel Organisms Using Transcriptomes and Low-Coverage Genomes: Improving Accuracy and Matrix Occupancy for Phylogenomics.” Molecular Biology and Evolution 31 (11). Oxford University Press: 3081–92.

